# Structure of *S. pombe* telomerase protein Pof8 C-terminal domain is an xRRM conserved among LARP7 proteins

**DOI:** 10.1101/739532

**Authors:** Ritwika Basu, Catherine D. Eichhorn, Ryan Cheng, Juli Feigon

**Affiliations:** Department of Chemistry and Biochemistry, P.O. Box 951569, University of California, Los Angeles, CA 90095-1569

**Keywords:** La protein, LARP, RRM, NMR, X-ray crystallography, telomerase, xRRM, RNA, 7SK

## Abstract

La related proteins group 7 (LARP7) are a class of RNA chaperones that bind the 3’ends of RNA and are constitutively associated with their specific target RNAs. In metazoa, Larp7 binds to the long non-coding 7SK RNA as a core component of the 7SK RNP, a major regulator of eukaryotic transcription. In ciliates, a LARP7 protein (p65 in *Tetrahymena*) is a core component of telomerase, an essential ribonucleoprotein complex that maintains the DNA length at eukaryotic chromosome ends. p65 is important for the ordered assembly of telomerase RNA (TER) with telomerase reverse transcriptase (TERT). Although a LARP7 as a telomerase holoenzyme component was initially thought to be specific to ciliate telomerases, *Schizosaccharomyces pombe* Pof8 was recently identified as a LARP7 protein and a core component of fission yeast telomerase essential for biogenesis. There is also evidence that human Larp7 associates with telomerase. LARP7 proteins have conserved N-terminal La motif and RRM1 (La module) and C-terminal RRM2 with specific RNA substrate recognition attributed to RRM2, first structurally characterized in p65 as an atypical RRM named xRRM. Here we present the X-ray crystal structure and NMR studies of *S. pombe* Pof8 RRM2. Sequence and structure comparison of Pof8 RRM2 to p65 and hLarp7 xRRMs reveals conserved features for RNA binding with the main variability in the length of the non-canonical helix α3. This study shows that Pof8 has conserved xRRM features, providing insight into TER recognition and the defining characteristics of the xRRM.

**Highlights:**

- The structure of the *S. pombe* LARP7 Pof8 C-terminal domain is an xRRM.
- Ciliates, human, and fission yeast contain LARP7 proteins with xRRMs involved in telomerase biogenesis.
- With three examples of xRRM structures, we refine the definition of xRRM.

## Introduction

The eukaryotic La protein and La-related protein (LARP) superfamily bind to diverse RNA targets and are involved in RNA processing and assembly (1–3). Genuine La protein recognizes the 3’UUU-OH terminus of most nascent RNA polymerase III transcripts to protect and stabilize them and in many cases also acts as a chaperone to fold the RNAs into functional complexes (4–10). LARPs bind to specific RNAs to function in the folding and biogenesis of their ribonucleoproteins (RNPs) (2). Based on evolutionary and domain organization conservation, they have been classified into four groups (LARP1, LARP4, LARP6 and LARP7), and the LARP7 family appears most closely related to genuine La protein (1–3).

All LARP7 proteins identified to date are components of 7SK (11,12) or telomerase RNPs (13–17). The 7SK RNP sequesters and inactivates the kinase activity of the positive elongation factor b (P-TEFb) to regulate the elongation phase of RNA Polymerase II (18,19) and has been identified in metazoa (20,21). Larp7 (here lowercase is used to distinguish the protein name in metazoa from the LARP7 family) binds to the long-noncoding 7SK RNA as a core component of the 7SK RNP and is required for 7SK RNP hierarchical assembly with P-TEFb and transcription regulation (11,12,22,23). Telomerase is an RNP that maintains telomeric DNA repeats at the ends of linear chromosomes (24,25) and has been identified in most eukaryotes (26,27). Telomerase is comprised of the telomerase RNA (TER) scaffold and template and the telomerase reverse transcriptase (TERT) enzyme, required together for catalytic activity, and additional proteins required for biogenesis and activity that form the holoenzyme *in vivo* (26,28,29). In the ciliates *Tetrahymena thermophila* and *Euplotes aediculatus*, the respective LARP7 protein p65 and p43 binds to TER and is required for its assembly with TERT (13,14,30–32). Recently, a LARP7 protein, Pof8 (also called Lar7), was identified in the fission yeast *Schizosaccharomyces pombe* (15–17). Pof8 binds to the *S. pombe* telomerase RNA (TER1) and recruits TER1 processing proteins Lsm2-8 to facilitate assembly of TER1 with telomerase reverse transcriptase (TERT), and is essential for telomerase biogenesis and function (15–17).

LARP7s have two structured domains – an N-terminal winged helix domain (La motif, LaM) followed by an atypical RNA recognition motif (RRM1) that together form a La module, and a C-terminal atypical RRM2 (2) (Fig 1A). The La module is a conserved feature of genuine La and LARPs that specifically recognizes and binds to the RNA 3’UUU-OH end (4,5,7), although the genuine La protein and LARP6 La modules sometimes bind alternate sequences (9,33,34). The La module of ciliate and metazoan LARP7s binds to the 3’UUU-OH terminus of their TER and 7SK cognate RNAs, respectively (23,35,36). For both *Tetrahymena* p65 and human Larp7, the C-terminal RRM2 is essential for specific RNA recognition and RNP assembly (22,31,32,37). High resolution structures of these domains in complex with RNA revealed an atypical mode of RNA binding divergent from that of canonical RRMs (38–40), and the domain was named xRRM for extended helical RRM (31,41,42). Compared to the canonical RRM, the xRRM lacks conserved single-strand RNA recognition sequences RNP1 and RNP2 on the β-sheet and has an additional C-terminal helix α3 that lies across this surface (Fig 1B,C) (40,41). These structures, together with multiple sequence alignment of LARP7s with predicted xRRMs, revealed several conserved features key for RNA recognition: a Y/W-X-D (RNP3) sequence on strand β2, a conserved R on strand β3, and charged/aromatic residues on the C-terminal end of helix α3 (Fig 1B,C) that together form an RNA binding surface on the side of the β-sheet rather than the surface and recognize a combination of base paired and unpaired nucleotides with high affinity and specificity (31,41,42). Human genuine La contains an atypical RRM with most of the features of an xRRM and has chaperone activity (43,44), but the RNA binding mode remains unknown.

**Figure 1.**
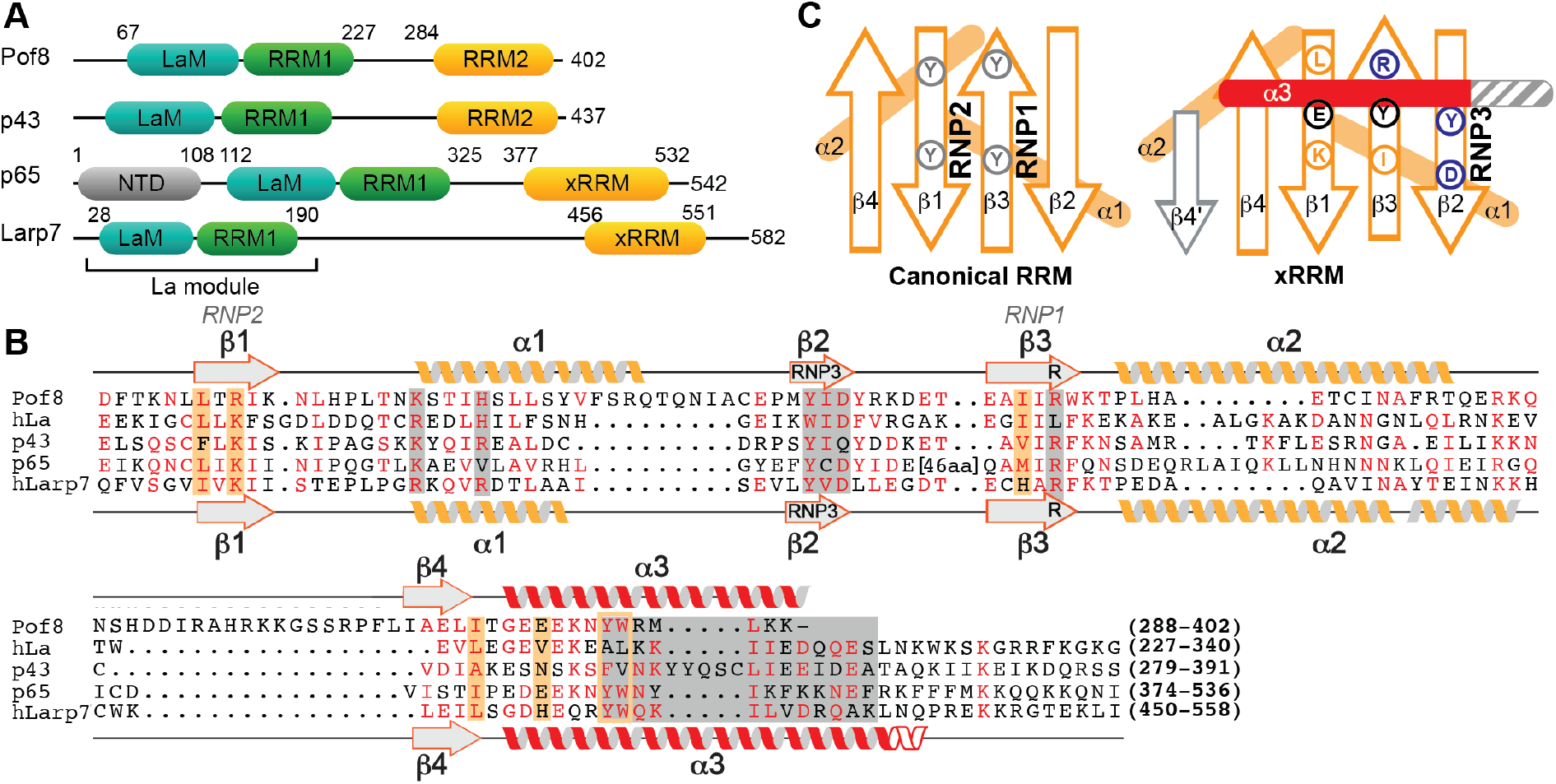
Domains and sequence alignments of LARP7 and La proteins. (A) Domain organization of LARP7s from yeast *S. pombe* (Pof8), ciliate *Euplotes aediculatus* (p43), ciliate *Tetrahymena thermophila* (p65), and human (Larp7). Numbers above the sequence schematic indicate known domain boundaries of the La motif (cyan), RRM1 (green) and xRRM (orange). (B) Sequence alignment of LARP7 and La proteins RRM2 whose structures have been determined, including Pof8 (this work), and *Euplotes* p43. The secondary structure elements of Pof8 and hLarp7 are shown as helices and sheets above and below the sequence, respectively. Conserved residues determined by this and previous work to contribute to RNA binding and helix α3–βsheet interaction are shaded gray and orange, respectively. (C) Cartoon comparing the conserved features of a canonical RRM (left) and xRRM (right), based in part on this work. Presence of β4’, length of helix α3 (shown in gray) and loop lengths vary.

The fission yeast LARP7 protein Pof8 was predicted to contain an xRRM at its C-terminus based on sequence similarity to *Tetrahymena* p65 and human Larp7 (15–17). Loss of Pof8 severely reduces telomerase activity *in vivo* and results in critically short telomeres, ultimately leading to uncapped chromosomes and chromosome end fusions (16). Truncation constructs deleting the putative RRM2 domain reduced TER1–TERT assembly, TER1 levels, and had a similar phenotype to Pof8 knock-down (16). Previously, association of a LARP7 protein with telomerase RNA was thought to be unique to ciliates, whose TER is an RNA polymerase III transcript with a native 3’UUU-OH terminus that binds the La module. Although this 3’-end sequence is absent in RNA polymerase II mRNA transcripts, the intron-encoded fission yeast TER1 is spliced resulting in a 3’UUU-OH terminus (45,46). However, the Lsm2-8 proteins bind to this region (47), ostensibly preventing La module interaction with the 3’terminus. The complete secondary structure of the 1213 nt TER1 has not been established and the binding site(s) for Pof8 is unknown. Although less well characterized, there is evidence that hLarp7 plays a role in human telomerase abundance and activity (48), suggesting that LARP7s may be broadly involved in telomerase function.

Here we present a 1.35 Å resolution crystal structure and solution NMR study of the C-terminal domain of Pof8, which reveals features consistent with an xRRM, despite having a shorter helix α3. We compare the structure with p65 and hLarp7 xRRM structures in the absence and presence of their cognate RNAs, as well as with genuine La RRM2, to refine the sequence and structural features of the xRRM class of atypical RRMs, and we propose the RNA binding mode in Pof8.

## Results and Discussion

### Crystal structure of the Pof8 C-terminal domain

Based on sequence homology and predicted secondary structure, a La module and RRM2 were predicted at the N- and C-termini, respectively (15–17) (Fig 1A). In particular, the C-terminal RRM2 domain of Pof8 has significant sequence homology to the xRRM domains in *Tetrahymena* p65 (31) and human Larp7 (49) (Fig 1B). Based on sequence alignment and known RRM topology (Fig 1C), a Pof8 construct containing residues 282-402 was cloned into a pET vector containing an N-terminal His_6_-SUMO fusion protein, and recombinant protein was expressed, purified, and screened by solution NMR spectroscopy to identify the presence of a folded domain. The Pof8 construct was crystallized, with crystals diffracting to 1.35 Å in space group P3121 (Table 1). The structure was determined by heavy atom (Hg) phasing using PCMBS soaked crystals. The electron density of the protein was visible up to 2σ, with weak density for residues 376-378 and no density was observed for N-terminal residues 282-287 or C-terminal residue 402 (Fig 2A). The crystal structure of Pof8 C-terminal domain revealed an atypical RRM with an overall β1-α1-β2-β3-α2-β4-α3 topology. A four-stranded antiparallel β-sheet consisting of β4 (aa 384-387), β1 (aa 294-298), β3 (aa 339-344) and β2 (aa 329-332) forms the front face with helices α1 (aa 306-320) and α2 (aa 347-359) packed on the back of the β-sheet to form the hydrophobic core and helix α3 (aa 391-401) on top of the β-sheet (Fig 2B,C). There are four loops: β1-α1 (aa 299-305), α1-β2 (aa 321-328), β2-β3 (aa 333-338), β3-α2 (aa 345-346), α2-β4 (aa 360-383), and β4-α3 (aa 388-390). Helix α3 (not present in canonical RRMs) is positioned orthogonal to the long axis of the β-strands, and lies across where the canonical RNP1 and RNP2 residues are normally found. Canonical RNP1 (<u>K/R</u>-G-<u>F/Y</u>-G/A-<u>F/Y</u>-I/L/V-X-<u>F/Y</u>) and RNP2 (I/L/V-<u>F/Y</u>-I/L/V-X/N/L) sequences on β3 and β1, respectively, are absent and there is a Y330–I331–D332 sequence on β2 (RNP3) and R343 on β3, consistent with an xRRM (Figs 1C, Fig 2D) (41).

**Table 1.** Crystallography statistics for *S. pombe* Pof8 xRRM

|  | Native | PCMBS soak |
| --- | --- | --- |
| **Data Collection** |  |  |
| Wavelength (Å) | 0.9791 | 0.97910 |
| Space Group | P31 2 1 | P31 2 1 |
| Cell dimensions |  |  |
| a, b, c (Å) | 57.36, 57.36, 70.3 | 56.01, 56.01, 65.29 |
| $\alpha$, $\beta$, $\gamma$ (°) | 90.0, 90.0, 120.0 | 90.0, 90.0, 120.0 |
| Resolution (Å) | 1.35 | 1.20 |
| R<sub>merge</sub> | 0.044 (0.68) | 0.055 (0.5) |
| R<sub>measure</sub> | 0.049 (0.78) | 0.054 (0.58) |
| R<sub>pim</sub> | 0.022 (0.36) | 0.019 (0.17) |
| I/$\sigma$ | 16.1 (1.95) | 17.1 (3.92) |
| Completeness (%) | 99.1 (98.6) | 96.2 (91.0) |
| No. of total reflections | 136156 | 162043 |
| No. of unique reflections | 29661 | 34507 |
| Multiplicity | 4.6 (4.3) | 10.0 (9.3) |
| Wilson B factor | 19.47 | 12.76 |
| CC<sub>1/2</sub> | 0.999 (0.83) | 0.998 (0.898) |
| CC* | 1 | 1 |
| **Refinement** |  |  |
| Resolution (Å) | 49.67-1.35 | 48.506-1.2 |
| No. of reflections | 29647 | 69061 (36157) |
| R<sub>work</sub>/R<sub>free</sub> | 0.1629/0.1940 | 0.2216/0.2313 |
| No. of atoms | 1019 | 901 |
| Protein | 936 | 888 |
| Nitrate | 4 | 4 |
| Water | 78 | 13 |
| Average B factors | 17 | 17 |
| r.m.s.d |  |  |
| Bond lengths (Å) | 0.005 | 0.007 |
| Bond angles (°) | 0.713 | 0.849 |
| Ramachandran plot |  |  |
| Favored (%) | 98.23 | 99.06 |
| Allowed (%) | 1.77 | 0.94 |
| Outliers (%) | 0.00 | 0.00 |

**Figure 2.**
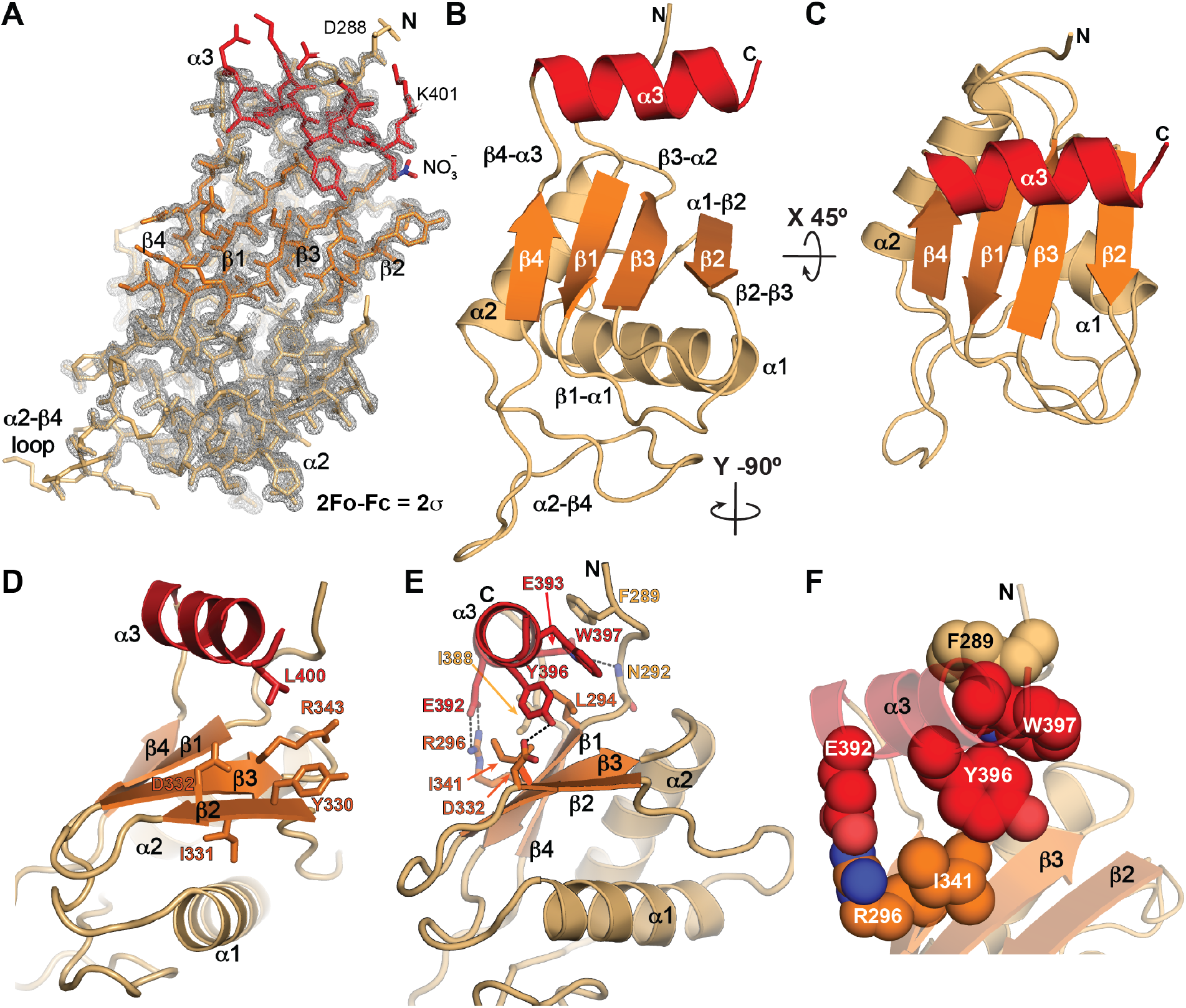
Crystal structure of Pof8 RRM2 at 1.35 Å. (A) 2Fo-Fc electron density map with crystal structure model shown in stick representation. The map is contoured at 2σ. (B, C) Two views of ribbon representation of the Pof8 xRRM (residues 288-402) crystal structure. The β-sheet is colored orange, helices α1 and α2 are tan, and helix α3 is red. (D) Ribbon representation with equivalent conserved residues involved in RNA binding in p65 and hLarp7 xRRMs shown as sticks. (E) Ribbon representation with conserved residues involved in stabilizing the α3–β-sheet interactions shown as sticks. (F) Ribbon representation with conserved residues involved in stabilizing the α3–β-sheet interactions shown as spacefill to highlight stacking interactions.

The Pof8 RRM2 helix α3 has three turns and is positioned through electrostatic and hydrophobic interactions with the β-sheet and the N-terminal residues (aa 289-294) (Fig 2E,F). There is a salt bridge between R296 (β1) and E392 (α3) side-chains, a hydrogen bond between the N292 backbone amide (N-tail) and the E393 side-chain (α3), and a hydrogen bond between D332 (β2) and Y396 (α3) side-chains. Residues L294 (β1), I341 (β3), I388 (β4-α3 loop), and Y396 (α3) form hydrophobic contacts at the β-sheet – helix α3 interface. The I341 (β3) side-chain stacks below the aromatic ring of Y396 (α3), and the W397 (α3) side-chain has π-π stacking with Y396 (α3) and F289 (N-tail) side-chains (Fig 2E,F). Overall, Pof8 RRM2 has all of the characteristic features of an xRRM except for the absence of residues beyond helix α3 that could extend it upon RNA binding.

### Conformational flexibility within the Pof8 RRM2

To investigate global and local dynamics of the Pof8 RRM2, the Pof8 construct used for crystal studies (residues 282-402) was also used for solution NMR studies. A 2D ^1^H-^15^N Heteronuclear Single Quantum Coherence (HSQC) spectrum of the backbone amides shows a well dispersed set of peaks indicative of a well-folded protein (Fig 3A). Backbone resonance assignments could be completed for the majority of the Pof8 RRM2, with the exception of α2-β4 loop residues Q350-L383 due to weak peak intensities caused by line broadening, indicative of conformational exchange. Close inspection of the 2D HSQC spectrum of the backbone amides revealed a set of uniformly low intensity additional peaks that appear to be due to peak doubling, likely caused by a second, lowly populated species in slow exchange with a major species (Fig 3A,B). 3D resonance assignment confirmed the identities of about half of these peaks. Peak doubling was only observed for the backbone amides. Nearly all of the doubled peaks where the second weak peak could be assigned were from residues at the β-sheet – helix α3 interface (T290, N292, K298, E339, I341, E386, E393, W397, E386) or the helix α3 C-terminus (R398, M399, L400, K401) (Fig 3C). Other weak peaks whose identities could be inferred from proximity to an assigned peak (e.g. L294, T295, R296) also correspond to residues at the β-sheet – helix α3 – N-tail interfaces (Fig 3A,C).

**Figure 3.**
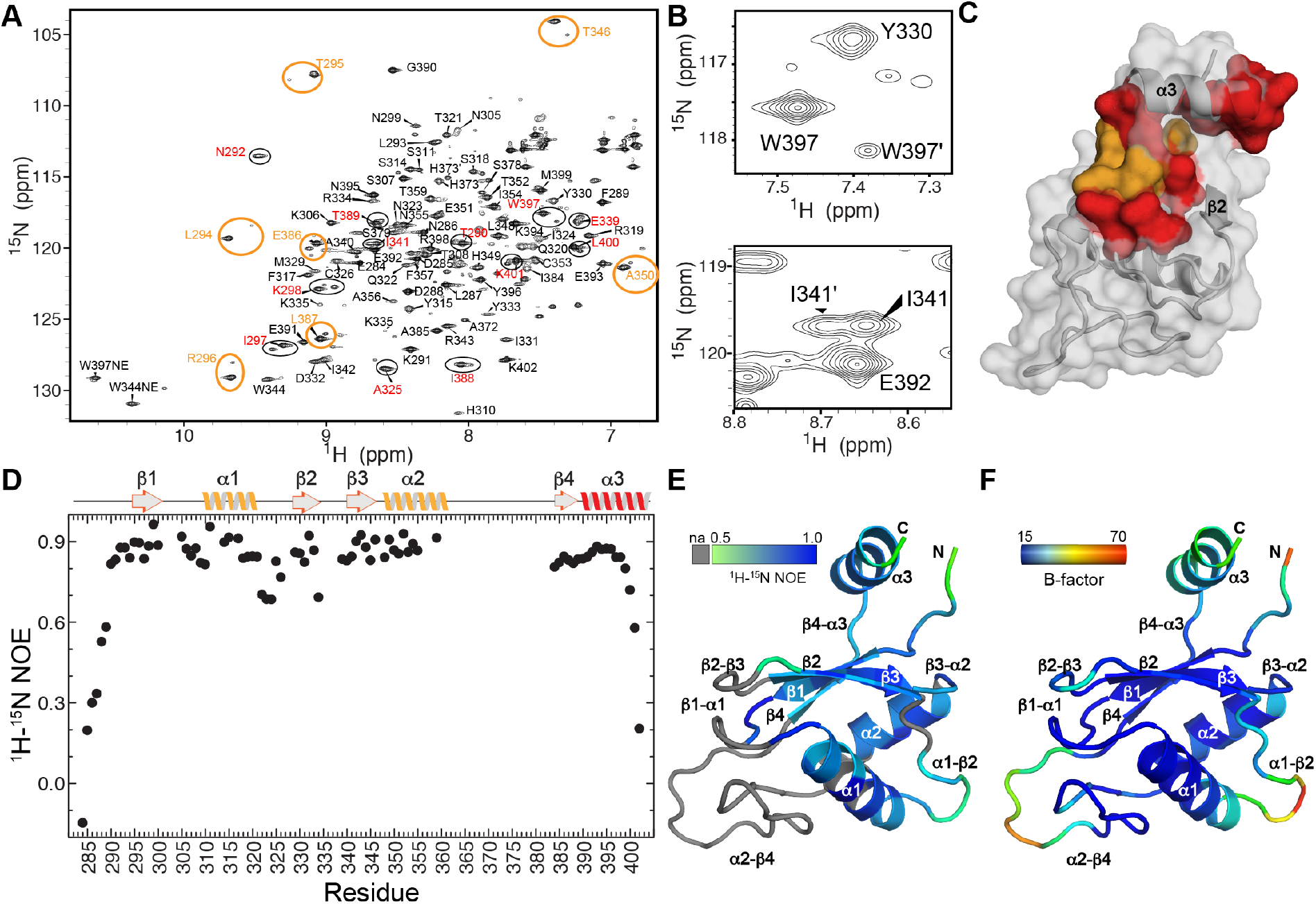
NMR characterization of Pof8 RRM2. (A) ^1^H-^15^N HSQC spectrum of Pof8 xRRM. Pairs of amide peaks that are doubled are circled; those for which the minor peak is assigned are labeled in red and inferred by proximity to major peak in orange. (B) Expanded regions from panel A showing peak doubling of amides W397 (helix α3) and I341 (β3). (C) Distribution of residues whose amides show peak doubling, mapped onto the structure. Assigned residues are red, residues with inferred assignments are orange. (D) Plot of heteronuclear NOE values vs residue number. Secondary structure elements are indicated above. (E) ^1^H-^15^N heteronuclear NOE values mapped onto a ribbon structure, with scale ranging from gray (na, not assigned), 0.5 (green) to 1.0 (blue). (F) Crystal structure B-factor mapped on a ribbon structure, scale ranging from 15.00 (blue) to 70.00 (red).

To determine the conformational flexibility in the major populated conformation of the Pof8 RRM2, we measured ^1^H-^15^N heteronuclear nuclear Overhauser effects (NOEs) (Fig 3D). Significantly reduced values are observed for the backbone amides of N-terminal residues E284-F289 and C-terminal residues 400-402, indicating that the N- and C-termini are highly flexible relative to the globular RRM fold. This data is consistent with the X-ray crystal electron density map that had missing density for N-terminal residues K282 to L287 and C-terminal residue K402. Reduced ^1^H-^15^N heteronuclear NOE values are also observed for residues Q322-C326 in the α1-β2 loop and residues R334-A340 (β2-β3 loop). ^1^H-^15^N heteronuclear NOE values could not be measured for the α2-β4 loop due to lack of resonance assignments (Fig 3D,E). Consistent with NMR data, the crystallographic B-factors indicate that the hydrophobic core is stable and that the N-terminus, α1-β2 loop, β2-β3 loop, α2-β4 loop, and helix α3 have higher B-factors (Fig 3F). In addition, although the α2-β4 loop has higher B-factors, there is clear albeit weak electron density for it (Fig 2A), while these residues could not be assigned due to conformational exchange. We attribute these differences to crystal contacts between the α2-β4 loop and another molecule in the crystal lattice. Together, these data indicate that for the Pof8 RRM2, the α2-β4 loop is flexible in solution and helix α3 appears to exist in two conformations: a major species where helix α3 is positioned on the β-sheet as in the crystal structure and a second minor species.

### Structural comparison of Pof8 with LARP7 and La protein RRM2s

The combination of helix α3, that lies across the β-sheet and over where the absent RNP2 and RNP1 would be, and conserved RNP3 sequence on the Pof8 RRM2 is unique to the xRRM class of atypical RRMs, and led us to hypothesize that the Pof8 RRM2 is an xRRM. The human genuine La protein also contains an atypical RRM2 that we previously predicted to be an xRRM (49). Below we compare the conserved structural and RNA recognition features for the existing examples of defined xRRMs (p65 and hLarp7) and predicted xRRMs (Pof8 and hLa).

Pof8, p65, hLarp7, and hLa RRM2 vary in the lengths of the loops between β strands and helices, presence or absence of a β4’, length of helix α3, and number of residues following α3 (Fig 4). For all, the beginning of helix α3 is enriched in acidic residues that face the β-sheet surface at β4 and β1, followed by aromatic and hydrophobic residues over β3 and β2, respectively (Fig 1B, Fig 4). Comparison of the three structures revealed several interactions between the β-sheet and helix α3 that are common to the structures of the LARP7 proteins. First, there is a salt bridge between a conserved (K/R) basic residue in the middle of β1 and an acidic residue in the first turn of helix α3, which is conserved in Pof8 (Fig 4A,E) and p65 (Fig 4B,F) but not present in hLarp7 (Fig 4C,G). Second, there is a hydrophobic patch between an I/L residue in β1, an I/L residue in the β4-α3 loop, and conserved Y-W residues in the second turn of helix α3 (Fig 4A-C,E,G). These hydrophobic contacts may stabilize the β4-α3 loop and help to orient helix α3 on the β-sheet. Third, there is a stacking interaction between a conserved hydrophobic/aromatic residue in β3 with the conserved Y in helix α3 (Fig 2F and Fig 4A-C,E-G). The contacts from β1 and β3 with residues on the first two turns of helix α3 anchor it on the β-sheet. Interestingly, the conserved residues in β1 and β3 that interact with helix α3 are located in the positions of RNP2 and RNP1, respectively, in the canonical RRM (Fig 1C) (50,51). It appears that while the xRRM lacks RNP1 and RNP2 sequences that interact with RNA, these sites on the β-sheet have adapted to interact with and stabilize helix α3 on the β-sheet surface.

**Figure 4.**
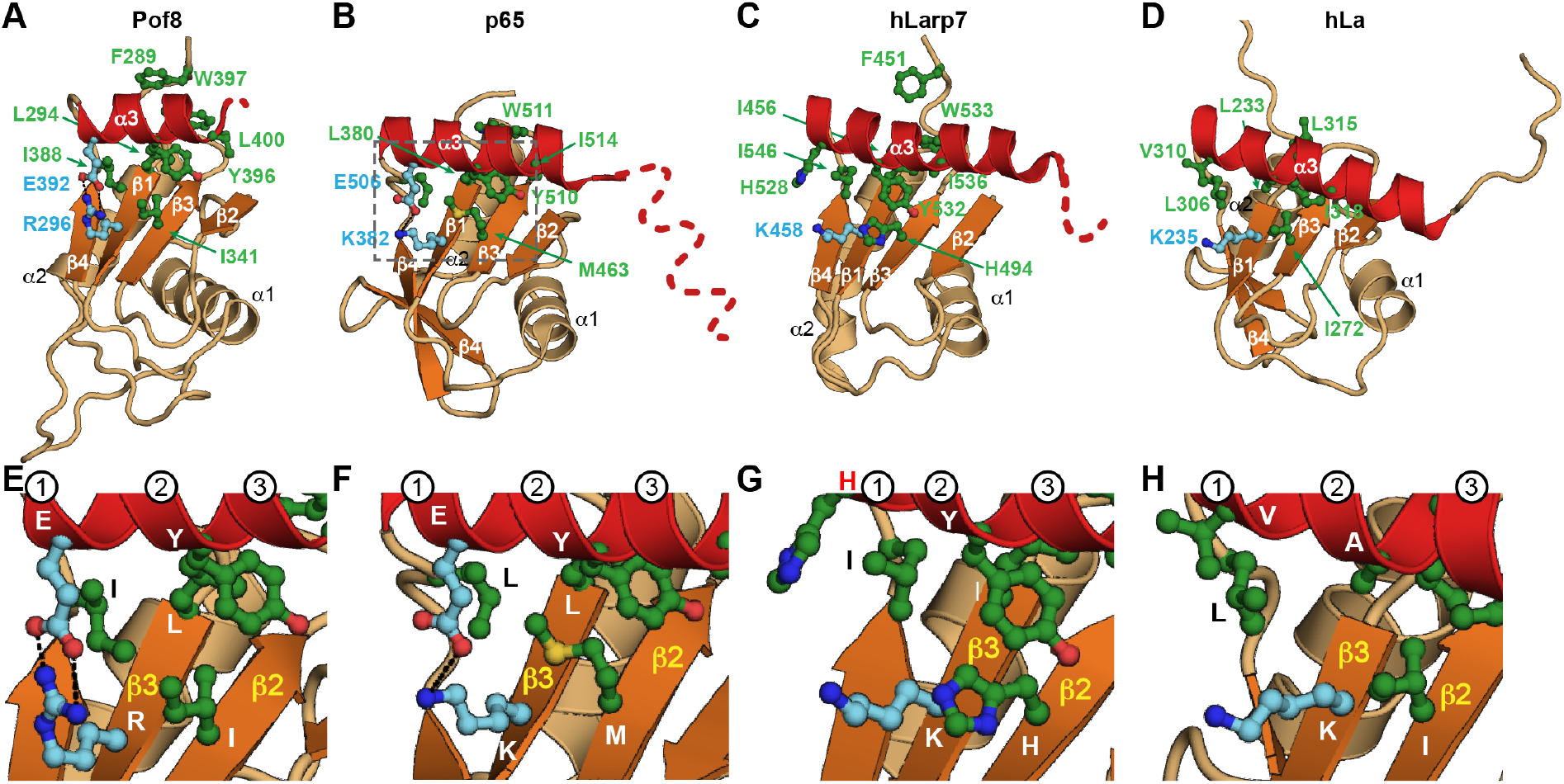
Comparison of structures of LARP7 and La protein xRRMs: helix α3–β sheet interactions. Ribbon representations of RRM2 from (A) *S. pombe* Pof8, crystal structure (this work), (B) *Tetrahymena* p65, crystal structure (PDB ID 4EYT) and (C) human Larp7, solution NMR structure (PDB ID 5KNW), (D) human La, solution NMR structure(PDB ID 1OWX). Dashed line indicates disordered residues missing in the density and/or that become helical on binding RNA. (E-H) Zoomed regions of (A-D) highlighting the conserved residues important for α3–β-sheet interactions. The first three turns of helix α3 are numbered. Secondary structure motifs are colored as in Fig 2. Residue side chains important for α3–βsheet interactions are shown as ball and stick and colored green (hydrophobic) and cyan (charged).

### RNA binding determinants of the xRRM

Based on the sequence and structural similarity of Pof8 to p65 and hLarp7 xRRMs (Fig 1B,C), comparison to structures of xRRM-RNA complexes (31,42), and effects of Pof8 amino acid substitutions and deletions on TER1 binding and abundance *in vivo* and telomere lengths (15,16), we can identify its putative RNA binding interface. Residues involved in RNA binding are shown as sticks on the ribbon model for p65 and hLarp7 (Fig 5A-C) (31,42). In structures of p65 and hLarp7 xRRMs in complex with their target RNAs, there is a conserved binding pocket for 2 or 3 nucleotides, respectively, including one Gua, that is formed by residues on the third and fourth turn of helix α3 and the β3-β2 strands (Fig 5E-G). The L on the third turn of helix α3 lies above the Gua, forming the ceiling of the binding pocket (Fig 5G). The RNP3 Y-X-D sequence on β2 and R on β3, previously identified as determinants of RNA binding (41), are identical among Pof8, p65, and hLarp7. For p65 and hLarp7, the RNP3 and R on β3 interact extensively with single-stranded RNA nucleotides, and alanine substitution of either Y (β2, RNP3) or R (β3) significantly impairs binding affinity to substrate RNA (31,42).

**Figure 5.**
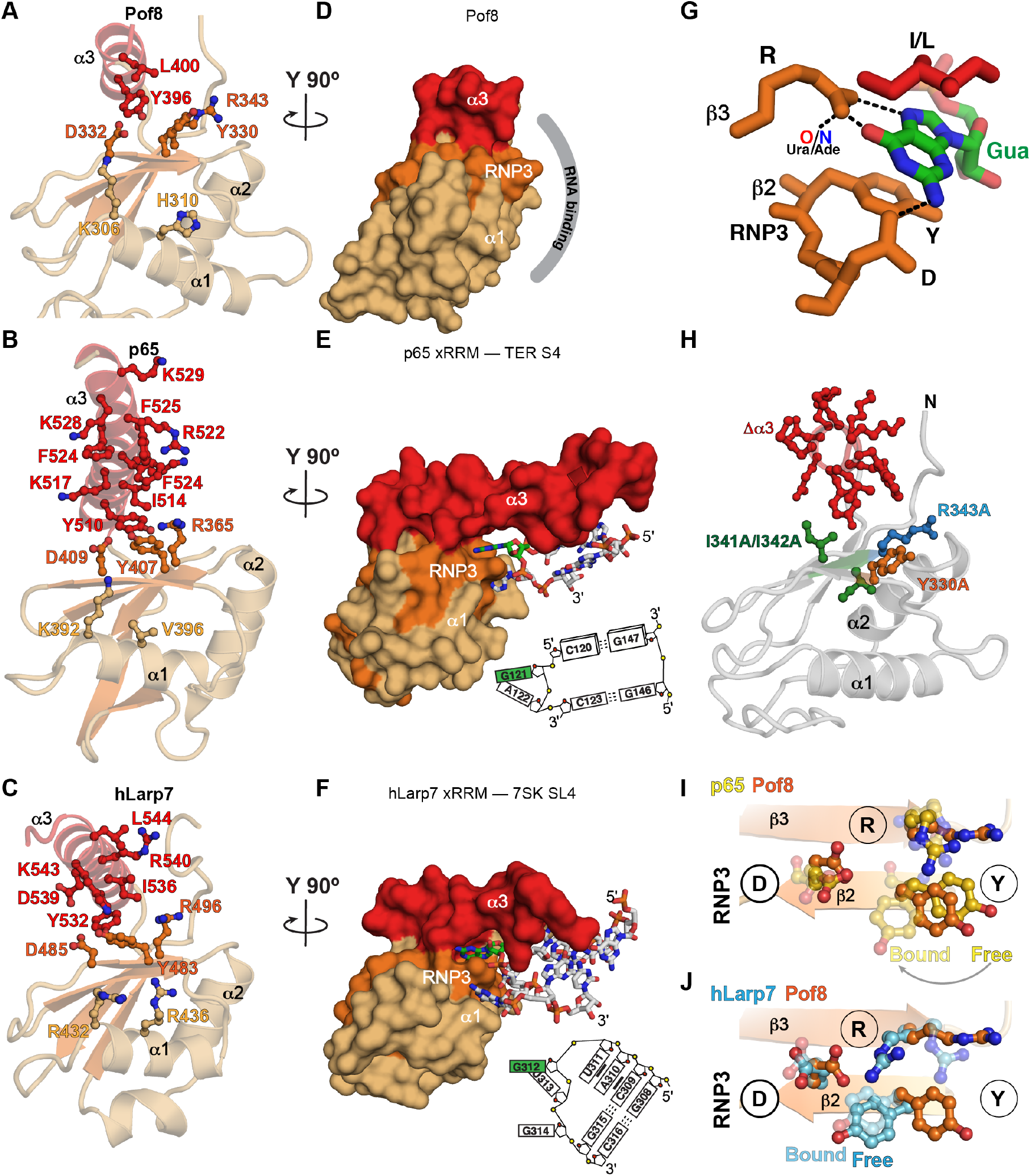
Comparison of structures of LARP7 and La protein xRRMs: RNA binding determinants. (A-C) Ribbon representations of crystal structures of (A) Pof8 xRRM (this work), (B) p65 xRRM bound to telomerase RNA stem 4 (PDB ID 4ERD), (C) hLarp7 xRRM bound to 7SK stem-loop 4 (PDB ID 6D12). Side chains that interact (B, C) or are predicted to interact (A) with RNA are shown as sticks; (D-F) Surface representations of (D) Pof8 xRRM (this work), (E) p65 xRRM bound to telomerase RNA stem 4 (PDB ID 4ERD), (F) hLarp7 xRRM bound to 7SK stem-loop 4 (PDB ID 6D12), rotated 90° from (A-C). The proposed RNA interacting surface of Pof8 is shown by the gray arc, and for p65 and hLarp7 the interacting RNA nucleotides are shown as sticks, with Gua in green, and an RNA schematic is shown at right; (G) Stick representation of Gua recognition in the xRRM binding pocket. The interaction between β3 R and a Ura O4 (e.g. hLarp7) or Ade N7 (e.g. p65) is also indicated; (H) Ribbon representation of Pof8 with residues whose substitution to alanine or deletion affect RNA binding in vitro or TER abundance and telomere length in vivo (15–17) shown as sticks. Δα3 is deletion of the entire helix. I341A/I342A is a double substitution. (I-J) Ribbon and stick representation illustrating position of RNP3 Y on (I) RNA free and bound p65 xRRM (gold) and RNA free Pof8 xRRM (orange) and (J) RNA free and bound hLarp7 xRRM (argon blue) and RNA free Pof8 xRRM (orange).

Figure **5H** maps mutations and deletions that affect Pof8 function *in vivo* onto the crystal structure of Pof8. Deletion of helix α3 (residues 390-402) reduced TER1 levels in a similar manner as Pof8 deletion, and caused telomere shortening (16) and substituting I341A-I342A, which would remove the hydrophobic contact between I341 (β3) and Y396 (α3), also resulted in shortened telomeres, reduced TER1 levels, and reduced co-immunoprecipitation of Pof8 with TER1 (15). These results indicate that helix α3 is essential for RNA binding and stability *in vivo* and are consistent with a requirement for stable positioning of helix α3 to form the RNA binding pocket. Single residue substitutions Y330A (RNP3) and R343A (β3) had similarly deleterious effects *in vivo* compared to deletion of helix α3, deletion of the RRM, or full-length knock-down (16), consistent with their predicted importance for nucleotide recognition in the putative Pof8 RNA binding pocket (Fig 5D).

In p65 and hLarp7, the binding pocket recognizes two or three nucleotides, respectively, that insert into the binding pocket between helix α3 and β3-β2, one of which is a Gua (Fig 5G). The conserved R on β3 has two hydrogen bonds to the Hoogsteen edge of Gua while the conserved D on RNP3 hydrogen bonds to the Watson-Crick face. The Gua is further stabilized in the binding pocket by stacking interactions with helix α3 conserved I (I/L) above and conserved RNP3 Y residue below. The β3 R also hydrogen bonds to AdeN7 (p65) or UraO4 (hLarp7) (Fig 5G). In p65, the RNP3 Y undergoes a large change in position between the RNA-free and RNA-bound states (Fig 5H), while in hLarp7 the position of RNP3 Y is already near to the bound state position prior to RNA binding (Fig 5I). For the RNA-free Pof8 structure reported here, the position of RNP3 Y is close to that of free p65 (Fig 5H,I). In p65 and hLarp7, the longer helix α3 also binds to two or one base pairs, respectively. In the case of p65 the long helix inserts between two stems on either side of the Gua-Ade bulge residues recognized in the binding pocket described above, causing a large bend between helices, while for hLarp7 a base pair at the top of the hairpin loop is recognized (Fig 5E,F). We speculate that in Pof8, the two conserved lysines at the end of helix α3 might hydrogen bond to a base pair. Based on the above analysis, we conclude that Pof8 has a binding pocket for a Gua (or possibly a Ura) and at least one other nucleotide analogous to that for hLarp7 and p65.

Finally, we note that in Pof8, p65, hLarp7, and hLa, helix α1 contains a conserved basic (K/R) residue (Fig 1B) that in hLarp7 interacts with RNA (Fig 5F) (42). A similar interaction may be present in the p65–RNA complex, but is not definitive since the sequence in this position in the crystal structure is not native (31). We propose that this interaction is common to the LARP7 proteins, and may provide further RNA binding affinity and/or specificity. Based on our analysis of the structures, sequence, and mutagenesis data, we conclude that the Pof8 RRM2 is an xRRM, as discussed further below.

### Comparison of LARP7 xRRMs to La RRM2

Human La protein RRM2 has known chaperone activity and has broad RNA substrate recognition (2,10). La protein generally binds RNA polymerase III transcripts temporally after transcription, but does not remain associated with the RNA, in contrast to LARP7s. There is an extensive hydrophobic interface and helix α3 lies closer to the β-sheet. hLa has an RNP3 (W261-I262-D263) characteristic of xRRMs, but the R on β3 that is conserved among LARP7s is replaced by an L (Fig 1B). This R contributes to nucleotide specificity of LARP7 xRRMs through hydrogen bonding to bases, and its replacement in hLa protein with an L might explain the lower binding affinity and lack of specificity of hLa RRM2 vs LARP7 RRMs (52,53). As there are no structures to date of hLa RRM2 in complex with RNA, the binding mode is unknown. Based on the analysis above, we propose that hLa RRM2 is an xRRM with reduced specificity in accordance with its function in binding multiple substrates.

### Conclusions

Pof8 and p65 are constitutive components of *S. pombe* and *Tetrahymena* telomerase, required for biogenesis and assembly of TER with TERT. The structure of Pof8 RRM2 reported here is the third example from the LARP7 family, and provides new insights into RNA binding by these domains and those of the related La proteins. The xRRM was first structurally characterized in p65 in the absence and presence of its RNA target (31), and subsequently in human Larp7 (42,49). Comparing the three LARP7 RRM2 structures shows that they share all the features of an xRRM first defined for p65 except for the length and sequence of helix α3that extends beyond the β-sheet, which are strikingly variable (Fig 4). Although in the absence of RNA the p65 helix α3 is a similar length to that of Pof8, upon RNA binding the helix is extended from four to eight turns (Fig 4B). In free hLarp7 α3 has 5 turns and is extended by an additional turn on binding RNA. For both p65 and hLarp7 xRRMs, truncation of these extra turns of helix α3 resulted in significantly reduced binding affinity to substrate RNA (31,49), indicating that the region of helix α3 that extends beyond the β-sheet contributes to high affinity binding.

The sequence differences in helix α3 between p65 and hLarp7 appear to aid in substrate discrimination; the aromatic residues in p65 helix α3 insert orthogonal to the helical axis in the TER SL4 major groove to induce a sharp bend in the RNA GA bulge, while the basic residues in the hLarp7 helix α3 interact with and insert parallel to the major groove at the RNA apical loop (31,42). The Pof8 α3 is three turns long and ends at the protein C-terminus with two consecutive lysines, the last of which is disordered. It seems likely that the short Pof8 helix α3 will result in a weaker binding affinity to the xRRMs of p65 (30 nM) (31) and hLarp7 (100 nM) (42,49) to their cognate substrates. Additional affinity may be provided by interactions between conserved residues on α1 and RNA. In summary, this work has defined the structure of the recently discovered yeast telomerase holoenzyme protein Pof8 C-terminal domain and provided a more definitive description of the xRRM and its conserved versus variable determinants of RNA specificity and affinity.

## Methods

### Protein expression and purification

The gene encoding Pof8 RRM2 (residues 282-402) was codon optimized for *E. coli* and synthesized into a gBlock (Integrated DNA Technologies) and subsequently cloned into a pET-His_6_-SUMO vector using Gibson ligation (New England Biolabs) (54). The recombinant plasmid was subsequently transformed into *Escherichia coli* BL21 (DE3) cells (Agilent Technologies) for protein expression. Bacterial cultures were grown in LB media with 50 μg/ml kanamycin at 37°C until OD_600_ reached 0.6, then induced with a final concentration of 0.5 mM IPTG for 18-24 h at 18 °C. Cells were harvested, sonicated in resuspension buffer (10% glycerol, 50 mM Tris pH 7.5, 750 mM NaCl, 30 mM imidazole, 3 mM NaN_3_, 0.1% Triton-X 100, 1 mM 2-carboxyethyl phosphine (TCEP), sonicated and centrifuged at 17,000 rpm for 45 mins. Clarified cell lysate containing His_6_-tagged protein was loaded onto a Ni–Sepharose affinity column (HisTrap HP; GE Healthcare). The column was washed with wash buffer (5% glycerol, 50 mM Tris pH 7.5, 750 mM NaCl, 30 mM imidazole, 3 mM NaN_3_, 1 mM TCEP), then eluted with elution buffer (5% glycerol, 50 mM Tris pH 7.5, 750 mM NaCl, 300 mM imidazole, 3 mM NaN_3_, 1 mM TCEP) to release bound His_6_-SUMO-tagged Pof8 RRM2. The eluate was incubated with SUMO protease (expressed and purified in-house from a pET28a vector with an N-terminal His_6_ tag) (approximately 0.5 mg protease to 5-10 mg Pof8 RRM2) for 3-4 hours at ambient temperature while dialyzing against a buffer containing 20 mM Tris pH 7.5, 250 mM NaCl, 5 mM β-mercaptoethanol. The cleaved Pof8 RRM2 was purified by loading the cleavage reaction onto the Ni–Sepharose affinity column and collecting the flow-through. After further purification by size-exclusion chromatography (HiLoad 26/600 Superdex 75; GE Healthcare) in crystallization buffer (20 mM Tris, pH 7.5, 100 mM NaCl, 1 mM TCEP) or NMR buffer (20 mM NaPO4 pH 6.1, 50 mM KCl, 1 mM TCEP), protein peak fractions were measured, pooled and concentrated using 3 KDa cutoff Amicon filters (Millipore Sigma). Unlabeled protein was purified from cells grown in LB media (Fisher Scientific) and uniformly ^15^N- or ^15^N,^13^C-labeled proteins were purified from cells grown in M9 minimal media with ^15^N ammonium chloride and/or ^13^C D-glucose (Cambridge Isotope Labs) as the sole nitrogen and/or carbon source, respectively.

### NMR Spectroscopy

NMR samples were concentrated to 0.1-0.8 mM in NMR buffer plus 5% D_2_O. NMR experiments were performed at 298 K on AVANCE 800 MHz Bruker spectrometer equipped with HCN cryoprobe. Nearly complete backbone (N, H, C, Cα, Cβ, Hα, Hβ) assignments were obtained for all residues except residues in the α2-β4 loop (E361-L383), using standard triple resonance assignment experiments (55,56). Briefly, 3D CBCACONH, HNCACB, HNCA, HBHACONH, HNCO and HNCACO experiments from the Bruker experimental suite were acquired on Topspin 4.0.7 (Bruker) using non-uniform sampling with 25% sparsity and a poisson-gap sampling schedule (57). The experiments were processed with Topspin 4.0.7 and analyzed using NMRFAM-SPARKY 1.414 (58) to assign backbone resonances (59,60). After partial manual assignment, the I-PINE web server was used to validate assignments and obtain additional assignments for residues in the β1-α1 loop L300 and T304, α1-β2 loop E327, β2-β3 loop K335-T338, β3-α2 loop K345, and α2 residue R358 (61). The ^15^N-3D NOESY experiment (noesyhsqcetf3gp3d) from the Bruker experimental suite was acquired on Topspin 4.0.7 (Bruker) with a mixing time of 120 ms to obtain NOE crosspeaks to further validate assignments. For I-PINE, atomic coordinates from the crystal structure, pre-assignments from manually assigned resonances, and peak lists from ^1^H-^15^N HSQC, ^1^H-^13^C HSQC, HNCA, HNCACB, CBCACONH, CCONH, HBHACONH, HNCO, HNCACO, and ^15^N-NOESY spectra were included as input. The ^1^H-^15^N heteronuclear NOE experiment (hsqcnoef3gpsi) from the Bruker experimental suite was acquired on Topspin 4.0.7 (Bruker), processed with NMRPipe (62), and analyzed with NMRFAM-SPARKY 1.414 (58) and Office Excel (Microsoft).

### Crystallization and data processing

The 2D ^1^H-^15^N HSQC of the Pof8 backbone amide resonances showed well-dispersed peaks, indicating a folded protein. The protein was concentrated to 25 mg/ml and crystallized in the hanging drop vapor diffusion method with reservoir solution 14% PEG 3350, 0.1 M NaNO_3_, 0.1 M Tris pH 8.5 and drops containing 1 μl of protein and 1 μl reservoir. Rod shaped crystals appeared in one day, were cryoprotected in reservoir solutions containing 35% PEG 3350 and flash frozen in liquid nitrogen. The native crystals diffracted to 1.35 Å. Crystals were soaked in 0.1 mM 4-chloro-mercuric-benzene-sulfonate (PCMBS) for 3-5 hours to obtain heavy atom (Hg) based experimental phases. All datasets were collected remotely at the Advanced Photon Source at Argonne National Labs, beamline 24-ID-C on a DECTRIS PILATUS 6M detector. The Matthews coefficient suggested that there was one molecule of Pof8 RRM2 in an asymmetric unit. XDS/XSCALE (63,64) was used to index, integrate and scale the data. Conservative resolution limits were applied based on I/σ, CC_1/2_ and R_sym_ values at the highest resolution shell. SAD phasing was performed by SHELXC/D/E (65) and HKL2MAP (66). Four Hg sites were identified by SHELXD. SHELXE was used to assign the handedness of the model, produce initial phases and solvent flattening. Phases were further improved with SHARP (67), and an initial model was generated with AUTOSHARP (68) and BUCCANEER (69). Native crystal structure was solved by Molecular Replacement using the Hg-model. The model was initially refined using Phenix version 1.13.2998 (70) and Coot (71), with final refinement performed using PHENIX with TLS refinement (72).

## Data availability

Atomic coordinates and structure factors have been deposited in the Protein Data Bank with the following accession codes: XXXX. NMR chemical shift assignments have been deposited in the BioMagnetic Resonance Bank with the following accession code XXX. All other data generated or analyzed in this study are included in the published article (and its supplementary information files) or are available from the corresponding author upon reasonable request.

## Acknowledgements

This work was supported by NSF grant MCB1517625 and NIH grant R35GM131901 to J.F. The authors acknowledge NMR equipment grants NIH S10OD016336 and S10OD025073 and DOE grant DE-FC0202ER63421 for partial support of NMR and X-ray core facilities. The authors thank M. Capel, K. Rajashankar, N. Sukumar, F. Murphy, I. Kourinov and J. Schuermann of Northeastern Collaborative Access Team (NE-CAT) beamline ID-24 at the Advanced Photon Source (APS) of Argonne National Laboratory, which are supported by NIH grants P41 RR015301 and P41 GM103403. Use of the APS is supported by DOE under Contract DE-AC02-06CH11357. The authors thank Dr. Lukas Sušac and Dr. Robert Peterson for help in the early stages of this work.

## Author Contributions

R.B. purified, crystallized, solved and analyzed the crystal structure of Pof8 xRRM; C.D.E. collected and analyzed NMR data and the structure; R.C. cloned, expressed, and purified Pof8 xRRM and collected NMR data; J.F. analyzed the structure and supervised the project; all authors contributed to manuscript writing, editing, and figure preparation.

